# Mechanism-Based Inhibition of Histone Deacetylase 6 by a Selenocyanate is Subject to Redox Modulation

**DOI:** 10.1101/2025.01.04.631333

**Authors:** Juana Goulart Stollmaier, B. Abigail R. Czarnecki, David W. Christianson

## Abstract

Organoselenocyanates have attracted considerable attention in recent years due to their therapeutic potential and versatility in medicinal chemistry. Here, we report on the mechanism of inhibition by 5-phenylcarbamoylpentyl selenocyanide (SelSA-2), an analogue of the well-characterized histone deacetylase inhibitor suberoylanilide hydroxamic acid (SAHA, a.k.a. Vorinostat). We show that histone deacetylases 6 and 10 can promote selenocyanate hydrolysis to generate a selenolate anion, and we explore the redox chemistry of selenium as it modulates inhibitory activity through reversible formation of the diselenide. The 2.15 Å-resolution crystal structure of histone deacetylase 6 cocrystallized with SelSA-2 conclusively demonstrates that it is not the selenocyanate, but instead the selenolate anion, that is the active pharmacophore responsible for enzyme inhibition.

---

The zinc-dependent histone deacetylases (HDACs 1–11)^1^ catalyze diverse amide hydrolysis reactions in all forms of life.^2-6^ These reactions most often comprise the deacetylation of lysine residues in protein substrates, but also include the deacylation of diverse lysine amides. For example, HDAC3 catalyzes lysine delactoylation,^7^ HDAC6 catalyzes lysine decrotonylation,^8^ HDAC6 and HDAC8 catalyze lysine depyruvoylation,^7^ and HDAC11 catalyzes lysine-fatty acid deacylation.^9-11^ Moreover, HDAC10 catalyzes the hydrolysis of a small molecule instead of a protein substrate, *N*^8^-acetylspermidine,^12,13^ so the HDAC substrate pool is vast and ranges from proteins to small molecules. Of note, the name “histone deacetylase” belies the chemical and biological diversity of this enzyme family, but this name is generally retained to reflect its historical roots.^14-16^

The HDACs are validated targets for the design and development of inhibitors that serve as cancer chemotherapy drugs. Suberoylanilide hydroxamic acid (SAHA, formulated as Vorinostat (Figure 1A)) was the first HDAC inhibitor approved for clinical use by the US Food and Drug Administration (FDA).^17,18^ Most recently, another hydroxamic acid-based inhibitor, Givinostat, was approved for the treatment of Duchenne muscular dystrophy, which is the first non-cancer indication approved for clinical treatment by an HDAC inhibitor.^19^

**Figure 1.**
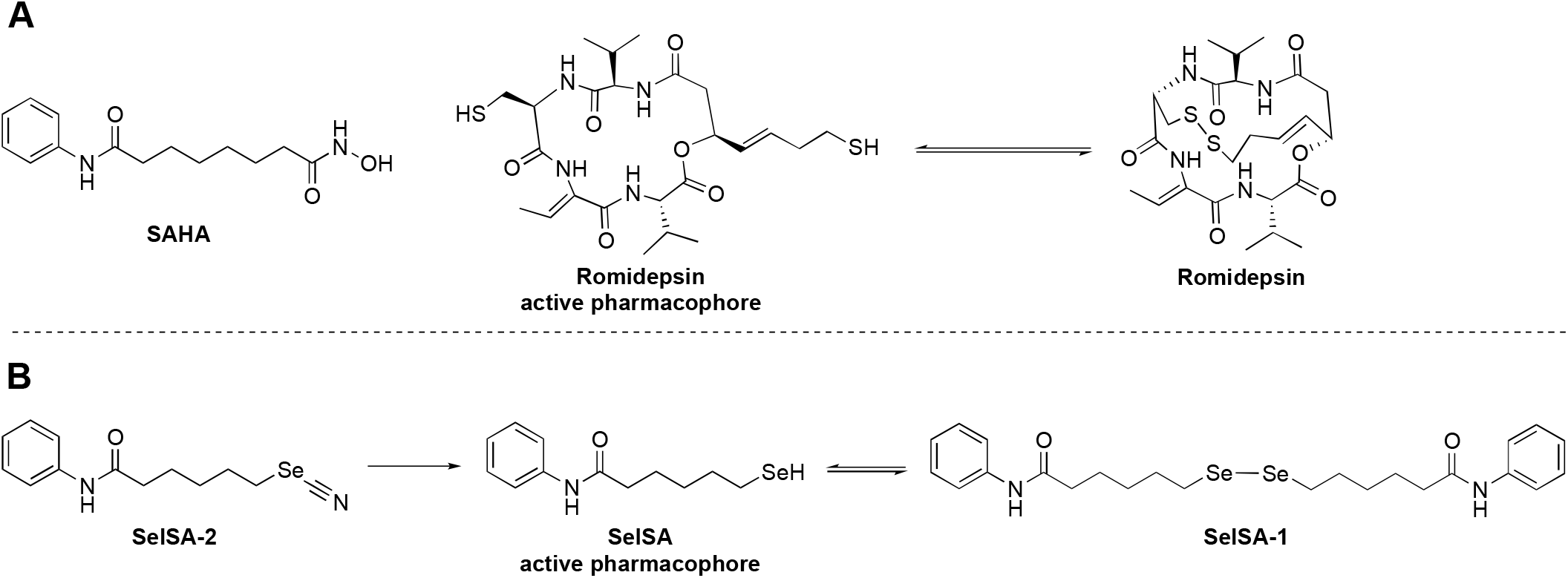
(A) Structures of selected FDA-approved HDAC inhibitors. Reduction of the disulfide linkage in Romidepsin yields the active pharmacophore with two free thiol groups, one of which coordinates to the catalytic zinc ion. (B) Reduction or hydrolysis of 5-phenylcarbamoylpentyl selenocyanide (SelSA-2) yields the active pharmacophore SelSA, which can then form the diselenide SelSA-1 under nonreducing conditions. SelSA-1 yields two equivalents of SelSA under the reducing conditions of the cell. The [SelSA]/[Sel A-1] ratio can be modulated by the strength and/or concentration of reducing agents.

Three out of four of the HDAC inhibitors currently approved by the FDA for clinical use contain hydroxamic acid moieties that, upon ionization, chelate the catalytic zinc ion in the enzyme active site,^20,21^ as first revealed in the crystal structure of a bacterial HDAC homologue complexed with SAHA.^22^ However, the hydroxamic acid moiety is not an ideal zinc-binding group *in vivo* due to its susceptibility to degradative reactions that generate mutagenic hydroxylamine or isocyanate species.^23,24^ This chemical disadvantage motivates the search for alternative zinc-binding groups, such as the thiol group of the reduced form of Romidepsin (Figure 1A), a cyclic depsipeptide approved by the FDA for cancer chemotherapy.^20,21^ Upon coordination to zinc, a thiol group ionizes to form a thiolate anion, which is a particularly favorable zinc ligand due to the size, charge, and polarizability of the sulfur atom. Several examples of thiol-bearing inhibitors have been studied in HDAC complexes, including a Romidepsin analogue and macrocyclic peptides.^25-29^

Sulfur-zinc coordination interactions contribute significantly to the high affinity and potency of HDAC inhibitors. In the Periodic Table of the Elements, sulfur belongs to Group 16, also known as chalcogens. Notably, other chalcogens such as selenium similarly contribute to high affinity and potency when incorporated into an HDAC inhibitor. For example, the selenium-based analogues of SAHA, SelSA-1 and SelSA-2 (Figure 1B), were designed and developed for HDAC inhibition.^30^ The diselenide SelSA-1 is reduced under cellular conditions to release two molecules of the active form of the inhibitor, SelSA (Figure 1B), with a selenol group that can coordinate to the catalytic zinc ion as a selenate anion, just as Romidepsin has a thiol group that can coordinate to the catalytic zinc ion as a thiolate anion.^30,31^ Similarly, it is hypothesized that the selenocyanate SelSA-2 (IC_50_ = 8.9 nM against HDAC6) is reduced to unmask the selenolate anion for zinc coordination.^30^ This chemistry can also be achieved through hydrolysis, given that treatment of selenocyanates with hydroxide yields selenolates as well as diselenides.^32^ Alternatively, it is proposed that the intact selenocyanate group of SelSA-2 coordinates to the catalytic zinc ion through the Se and N atoms to yield pentavalent trigonal bipyramidal zinc coordination geometry.^33,34^ Thus, the mode of action of a selenocyanate inhibitor remains ambiguous.

To resolve this ambiguity, we cocrystallized zebrafish HDAC6 catalytic domain 2 (henceforth simply “HDAC6”)^35^ with selenocyanate SelSA-2 and determined the structure of the enzyme-inhibitor complex at 2.15 Å resolution. Complete experimental details are reported in the Supporting Information. Polder omit maps of the bound inhibitor clearly show the binding of SelSA (Figure 2, Figure S1). Thus, the active pharmacophore is the selenol SelSA and not the selenocyanate SelSA-2. Given that the pKa value for an alkylselenol is 5–7,^36^ and given that zinc coordination will further lower this pKa, the selenol group of SelSA is predominantly ionized as the selenolate anion.

**Figure 2.**
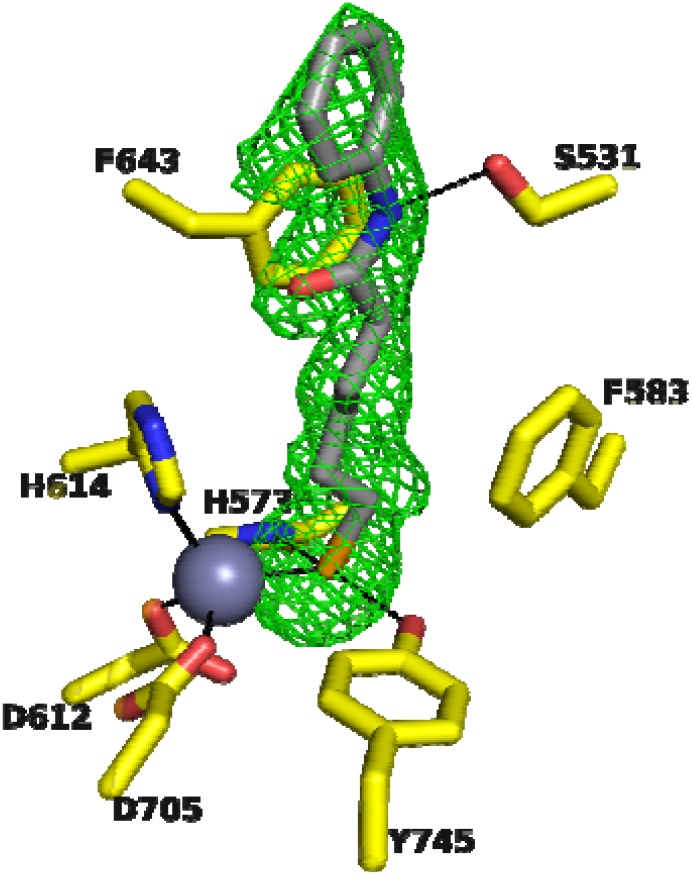
Polder omit map contoured at 3.0σ (green mesh) showing SelSA bound in the HDAC6 active site (monomer A in the crystal unit cell). Atomic color codes are as follows: C = yellow (HDAC6) or light gray (SelSA), Se = light orange, N = blue, O = red, Zn^2+^ = gray sphere; metal coordination interactions ≤ 2.5 Å and hydrogen bond interactions ≤ 3.4 Å are indicated by solid and dashed black lines, respectively.

The structure of the enzyme-inhibitor complex reveals that the selenolate anion of SelSA coordinates to zinc with a Se•••Zn^2+^ separation of 2.5 Å in both monomers. Additionally, the selenolate anion accepts hydrogen bonds from the catalytic tyrosine, Y745, and one of the catalytic histidine residues, H573. The benzamide NH donates a hydrogen bond to the side chain of S531, a residue unique to HDAC6 that confers selectivity for inhibitor binding through direct or water-mediated hydrogen bonds.^35,37,38^

The selenolate form of SelSA is thus established as the actual inhibitory species derived from SelSA-1 or SelSA-2, but how is the selenolate formed from the selenocyanate? Surprisingly, the enzyme can play a role in the chemical conversion of selenocyanate SelSA-2 into selenolate SelSA, which can then form diselenide SelSA-1 under nonreducing conditions.

In the absence of enzyme, redox chemistry governs the speciation of selenium, since incubation of SelSA-2 in size-exclusion (SEC) buffer at room temperature rapidly yields SelSA and SelSA-1 (Table S2). The SEC buffer contains the reducing agent tris(2-carboxyethyl)phosphine (TCEP), which promotes rapid reduction of the selenocyanate moiety of SelSA-2 to yield SelSA, which can then form SelSA-1. Notably, the [SelSA]/[SelSA-1] ratio is TCEP-dependent. Under atmospheric exposure *in vitro*, the reducing agent must be sufficiently concentrated to reduce both SelSA-2 and thence SelSA-1 to ensure that the active pharmacophore SelSA is stabilized. In SEC buffer lacking TCEP, SelSA-2 is stable and unreactive, with no conversion to SelSA or SelSA-1 even after 24 h. Thus, the concentration of the selenol pharmacophore in the absence of enzyme is regulated by two redox reactions – the initial reduction of the selenocyanate and the subsequent reduction of the diselenide.

The effect of reducing agents on the stabilization of SelSA is also observed during inhibitory activity assays performed with and without TCEP in the assay buffer (complete experimental details are reported in the Supporting Information, Figure S2). SelSA-2 does not inhibit HDAC6 in TCEP-free buffer, indicating that neither SelSA-1 nor SelSA-2 is the active pharmacophore. Inhibition of HDAC6 is observed only when SelSA-2 is incubated with TCEP, mostly due to the stabilization of SelSA in its equilibrium with SelSA-1.

With the effects of reducing agent thus established in the absence of enzyme, does the addition of enzyme influence the chemistry of selenium? Incubation of HDAC6 with SelSA-2 for 24 h in TCEP-free SEC buffer yields a 72%/28% mixture of SelSA-1/SelSA-2. Incubation of the related class IIb isozyme HDAC10 with SelSA-2 for 24 h in TCEP-free SEC buffer yields a 96%/4% mixture of SelSA-1/SelSA-2. These results suggest that the selenolate can also be generated slowly through selenocyanate hydrolysis in the presence of enzyme, which then leads to diselenide formation under nonreducing conditions. Cyanic acid is a co-product of this reaction, which is further hydrolyzed^39^ to form carbon dioxide and ammonium ion (Figure 3). To account for SelSA-1 formation under nonreducing or insufficiently-reducing conditions, enzyme-bound SelSA must dissociate into solution, where it can react with a second molecule of SelSA-2 to form the diselenide and a cyanide ion (Figure 3).

**Figure 3.**
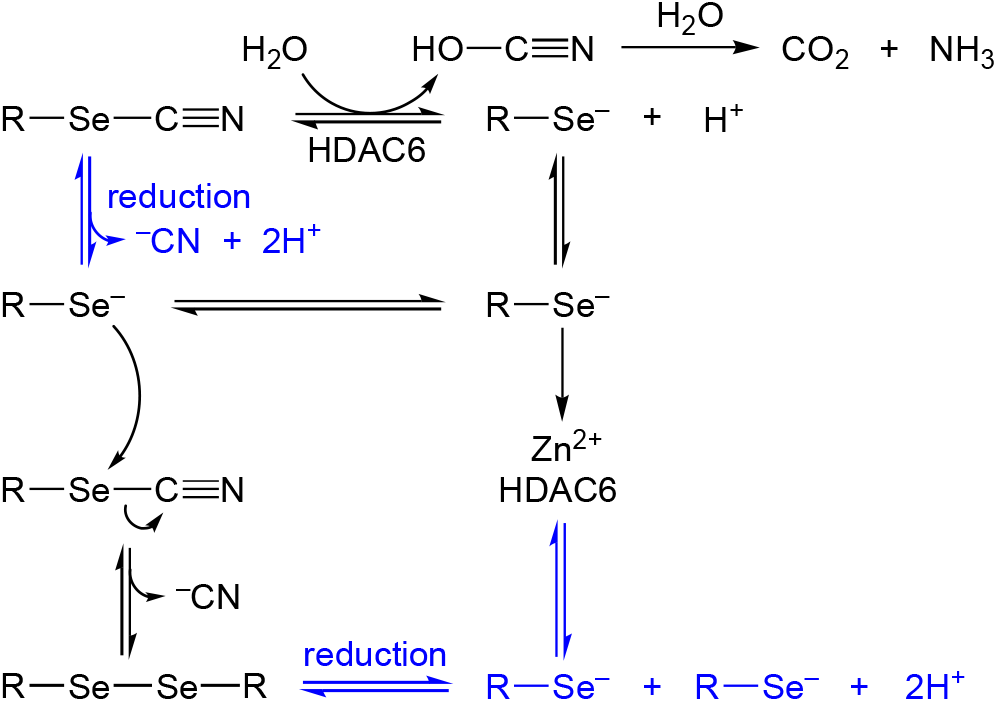
The selenocyanate moiety of SelSA-2 is converted to the selenolate anion SelSA by nonenzymic reduction or by enzyme-promoted hydrolysis, after which the selenolate anion coordinates to the catalytic zinc ion. The selenolate anion can also react with another selenocyanate to yield the diselenide under nonreducing conditions. Under sufficiently strong reducing conditions *in vitro* or *in vivo*, the selenolate anion is stabilized over the diselenide.

The above results implicate the zinc-bound water molecule in the HDAC6 and HDAC10 active sites as the nucleophile responsible for selenocyanate hydrolysis. This water molecule is regarded as an incipient hydroxide ion resulting from deprotonation by a histidine general base in catalysis. Correspondingly, when SelSA-2 is incubated with increasing concentrations of NaOH in SEC buffer lacking reducing agent or enzyme, complete conversion of SelSA-2 into SelSA-1 is observed at higher NaOH concentrations (Table S4). Similar chemistry has been previously observed for other selenocyanates hydrolyzed by NaOH or KOH in mixed aqueous-organic solutions.^32^ Therefore, we conclude that SelSA-2 is a mechanism-based inhibitor activated by the zinc-bound water molecule in the enzyme active site, with the stability of the resulting pharmacophore SelSA regulated by the strength of reducing conditions to the extent that SelSA is in equilibrium with SelSA-1.

Intriguingly, nucleophilic attack of zinc-bound water at the carbon-nitrogen triple bond of a selenocyanate is reminiscent of nucleophilic attack at the carbon-nitrogen double bond of an oxadiazole in the active site of HDAC6.^40^ In both examples, the reactivity of the zinc-bound water molecule is harnessed to convert a small molecule into a potent inhibitor. Such mechanism-based inhibitor activation represents a unique facet of diversity and versatility in the chemistry of HDAC inhibition.^41^

Organoselenium compounds are increasingly prominent in medicinal chemistry and drug design, especially in contexts where the rich redox chemistry or metal-binding properties of selenium can be harnessed.^42-45^ For example, diphenyldiselenide (PhSe)_2_ reduces reactive oxygen species in herpes simplex virus-infected mice,^46,47^ and phenylselenol (PhSeH) and its derivatives have been studied in complexes with human carbonic anhydrases that feature potent selenolate-zinc coordination.^48-50^ Because of the reactivity of the free selenol, it can be protected as a selenoester or selenocarbamate that undergoes enzyme-promoted hydrolysis to liberate an inhibitory selenol, which is then precisely positioned for zinc coordination.^48-50^

Here, we provide the first proof that a selenocyanate can similarly undergo enzyme-promoted hydrolysis to liberate an inhibitory selenol, which is then precisely positioned for zinc coordination as the selenolate anion in the HDAC6 active site. These results establish a critical foundation for further exploration of the biological and medicinal chemistry of selenium, as well as the redox regulation of this chemistry *in vitro* and *in vivo*.

## ASSOCIATED CONTENT

### Supporting Information

The Supporting Information is available free of charge on the ACS Publications website.

Complete materials and methods; Table S1, crystallographic data collection and refinement statistics; Table S2, redox speciation in the presence and absence of TCEP; Figure S1, stereoviews of electron density maps showing SelSA bound to monomers A and B in the unit cell; Figure S2, enzyme activity assay in the presence and absence of TCEP.

### Accession Code

The atomic coordinates and crystallographic structure factors for the HDAC6-SelSA complex have been deposited in the Protein Data Bank (www.rcsb.org) with accession code 9MQP.

## AUTHOR INFORMATION

**Authors**

**Juana Goulart Stollmaier** – Roy and Diana Vagelos Laboratories, Department of Chemistry, University of Pennsylvania, Philadelphia, Pennsylvania, 19104-6323, United States;

**Briana Abigail R. Czarnecki** – Roy and Diana Vagelos Laboratories, Department of Chemistry, University of Pennsylvania, Philadelphia, Pennsylvania, 19104-6323, United States;

## >Author Contributions

The research was performed and the manuscript was written through contributions of all authors. All authors have given approval to the final version of the manuscript.

## Notes

The authors declare no competing financial interests.

## ACKNOWLEDGMENT

This work is based on research conducted at beamline 17-ID-1 (AMX) of the National Synchrotron Light Source II, a DOE Office of Science User Facility operated for the DOE Office of Science by Brookhaven National Laboratory under Contract DE-SC0012704. The Center for BioMolecular Structure (CBMS) is primarily supported by the National Institutes of Health, NIGMS, through a Center Core P30 Grant (P30GM133893) and by the DOE Office of Biological and Environmental Research (KP1607011). We thank Alex Rodway for designing the TOC artwork.

## ABBREVIATIONS

BME: β-mercaptoethanol
DTT: dithiothreitol
FDA: Food and Drug Administration
HDAC: histone deacetylase
SAHA: suberoylanilide hydroxamic acid
SelSA: selenolate form of the inhibitor
SelSA-1: diselenide form of the inhibitor
SelSA-2: selenocyanate form of the inhibitor
TCEP: tris(2-carboxyethyl)phosphine.

## For TOC Use Only

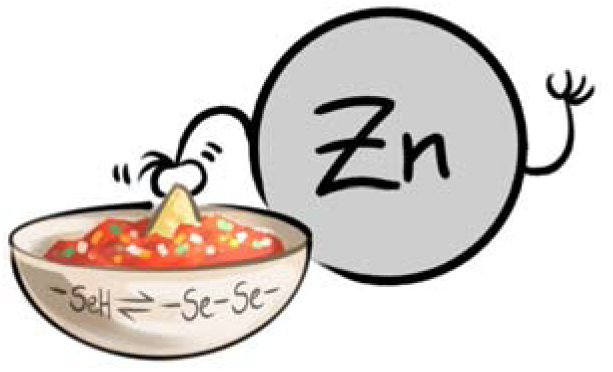

