## Supporting Information for "Mechanism-Based Inhibition of Histone Deacetylase 6 by a Selenocyanate is Subject to Redox Modulation"

Juana Goulart Stollmaier, Briana Abigail R. Czarnecki, and David W. Christianson*

Roy and Diana Vagelos Laboratories, Department of Chemistry, University of Pennsylvania, 231 South 34th Street, Philadelphia, PA 19104-6323 USA

### Materials and Methods

#### HDAC6 Expression and Purification

Catalytic domain 2 from *Danio rerio* (zebrafish) HDAC6 (henceforth simply "HDAC6") was expressed and purified as previously detailed,[1] with a small modification of 0 mM imidazole in buffer A.

#### HDAC10 Expression and Purification

HDAC10 from *D. rerio* containing the A24E and D94A substitutions to mimic the human enzyme (henceforth simply "HDAC10") was prepared and purified as previously described.[2]

#### Cocrystallization and Structure Determination of the HDAC6–SelSA Complex

A protein solution containing HDAC6 and SelSA-2 was prepared by adding 2 mM of SelSA-2 to a protein solution comprised of 10 mg/mL HDAC6 in size exclusion (SEC) buffer [50 mM HEPES (pH 7.5), 100 mM KCl, 5% glycerol (v/v), 1 mM TCEP] and equilibrated on ice for 2 h. The protein solution was filtered using 0.22 μm centrifuge filters. The enzyme-inhibitor complex was crystallized by the sitting-drop vapor diffusion method at 4 °C. A 200-nL drop of a freshly filtered protein solution was added to a 100-nL drop of precipitant buffer [0.09 M Bis-Tris (pH 6.0), 0.18 M potassium thiocyanate, 0.01 M cadmium chloride hydrate, 18% (w/v) PEG 3350], followed by a 25-nL drop of microseed (apo-HDAC6) solution. The sitting drop was equilibrated against 80 μL of precipitant buffer in the well reservoir of a 96-well crystallization plate using a Mosquito crystallization robot (TTP Labtech). Clusters of thin needle-like crystals formed in 5 days.

X-ray diffraction data from these crystals were collected on the NSLS-II AMX beamline at Brookhaven National Laboratory. Initial data processing was achieved using the autoPROC toolkit.[3]Data were indexed and integrated with XDS[4] and POINTLESS,[5] and scaled with AIMLESS[6] in the CCP4i2 program suite.[7] The initial electron density map was phased by molecular replacement using Phaser[8] with the atomic coordinates of unliganded catalytic domain 2 of HDAC6 (PDB 5EEM) used as a search probe.[9] The protein model was manually adjusted as needed using Coot[10] and refined in Phenix.[11] The inhibitor and water molecules were fit to the electron density map in the later stage of refinement. Notably, the electron density map clearly indicated the binding of SelSA rather than SelSA-2. Validation of the final model was performed with MolProbity.[12] Data collection and refinement statistics are recorded in Table S1.

**Table S1.** Crystallographic Data Collection and Refinement Statistics for the HDAC6–SelSA Complex

| | |
|---|---|
| **Unit Cell** | |
| Space group | *P*1 |
| *a, b, c* (Å) | 48.17, 54.19, 74.35 |
| $\alpha, \beta, \gamma$ (deg) | 73, 90, 83 |
| **Data Collection** | |
| Laboratory, beamline | NSLS-II<br>17-ID-1 AMX |
| Detector | EIGER 9M |
| Resolution (Å) | 2.15 |
| Total/unique no. of reflections | 79,465/38,156 |
| $R_{merge}$[a,b] | 0.224 (0.565) |
| $R_{pim}$[a,c] | 0.194 (0.495) |
| $CC_{1/2}$[a,d] | 0.765 (0.624) |
| $I/\sigma(I)$[a] | 2.2 (1.0) |
| Redundancy[a] | 2.1 (2.0) |
| Completeness (%)[a] | 98.3 (97.5) |
| **Refinement** | |
| Reflections used in refinement/test set | 38,151/1,999 |
| $R_{work}$[a,e] | 0.225 (0.285) |
| $R_{free}$[a,e] | 0.239 (0.355) |
| No. of protein chains | 2 |
| No. of nonhydrogen atoms | 5,703 |
| Protein | 5,455 |
| Ligand | 44 |
| Solvent | 204 |
| Average *B* factor ($Å^2$) | 14 |
| Protein | 14 |
| Ligand | 17 |
| Solvent | 16 |
| **Root-mean-square deviation from ideal geometry** | |
| Bonds (Å) | 0.004 |
| Angles (deg) | 0.8 |
| **Ramachandran plot**[f] | |
| Favored (%) | 96.40 |
| Allowed (%) | 3.45 |
| Outliers (%) | 0.14 |
| Molprobity score[f] | 1.34 |
| PDB accession code | 9MQP |

[a]Values in parentheses refer to the highest-resolution shell of data. [b]$R_{merge} = \sum_h \sum_i |I_{h,i} - \langle I \rangle_h| / \sum_h \sum_i I_{h,i}$, where $\langle I \rangle_h$ is the average intensity calculated for reflection *h* from *i* replicate measurements. [c]$R_{p.i.m.} = (\sum_h (1/(N-1))^{1/2} \sum_i |I_{h,i} - \langle I \rangle_h|) / \sum_h \sum_i I_{h,i}$, where N is the number of reflections and $\langle I \rangle_h$ is the average intensity calculated for reflection *h* from replicate measurements. [d]Pearson correlation coefficient between random half-datasets. [e]$R_{work} = \sum||F_o| - |F_c|| / \sum|F_o|$ for reflections contained in the working set. $|F_o|$ and $|F_c|$ are the observed and calculated structure factor amplitudes, respectively. $R_{free}$ is calculated using the same expression for reflections contained in the test set held aside during refinement. [f]Calculated with MolProbity.

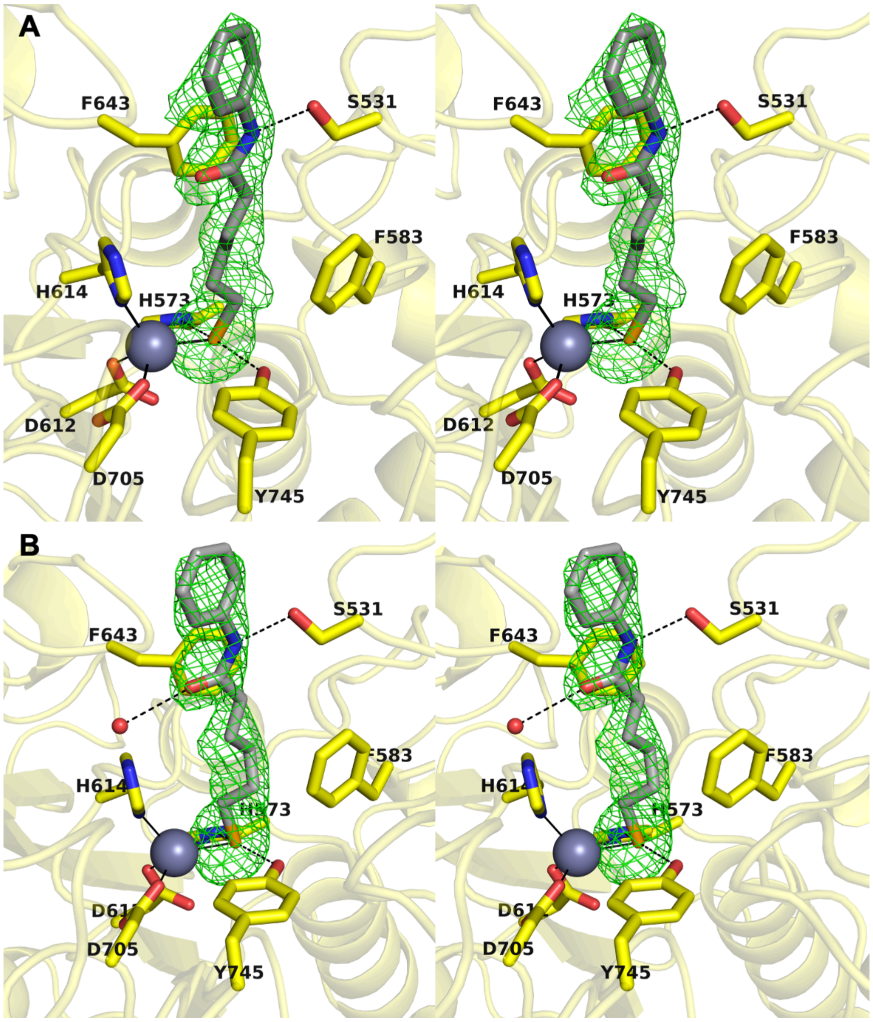


**Figure S1.** Stereoviews of Polder omit maps contoured at 3.0σ (green mesh) showing SelSA bound in Chain A (A) and Chain B (B) of the HDAC6 active site. Atomic color codes are as follows: C = yellow (HDAC6, both chains) or light gray (SelSA, Chain A and Chain B), Se = light orange, N = blue, O = red, $Zn^{2+}$ = gray sphere; metal coordination interactions ≤ 2.5 Å and hydrogen bond interactions ≤ 3.4 Å are indicated by solid and dashed black lines, respectively. Selected active site residues are shown as stick figures and labeled.

**Inhibitor speciation assays using ultra-performance liquid chromatography-mass spectrometry (UPLC-MS) analysis**

To determine the effects of reducing agent and enzyme on inhibitor speciation, i.e., whether the inhibitor was stabilized as SelSA-2, SelSA, or SelSA-1, UPLC-MS was employed. To ascertain inhibitor speciation in buffer solutions without enzyme, varied concentrations of SelSA-2 were incubated in size exclusion buffer (as formulated for HDAC6 or HDAC10) for 30 min at room temperature, total volume 100 µL. After incubation, samples were further diluted with 100 µL methanol. To ascertain inhibitor speciation in the presence of enzyme, 50 µM HDAC10 or HDAC6 in corresponding SEC buffer was incubated with 100 µM of SelSA-2, total volume 100 µL. For experiments performed in SEC buffer without TCEP, 50 µM HDAC6 or HDAC10 was dialyzed in corresponding SEC buffer without TCEP for 1 h prior to the addition of inhibitor. Following incubation for 18 h at room temperature, protein was precipitated using methanol (100 µL) followed by filtering through a 22-µm GV Durapore filter.

A 2-µL aliquot of each mixture was injected over a C18 reverse-phase column on a Waters Acquity UPLC-MS using a 2-min gradient of 95:5 $H_2O$:MeCN to 5:95 $H_2O$:MeCN. Mass spectra were analyzed using MestreNova (Mestrelab Research). Measured percentages of SelSA-2, SelSA and SelSA-1 are recorded in Table S2.

**Table S2.** Inhibitor Speciation in the Presence and Absence of TCEP

| **Entry** | **Buffer type** | **[TCEP] (mM)** | **[SelSA-2] (µM)** | **SelSA-2 (%)** | **SelSA (%)** | **SelSA-1 (%)** | **Alcohol byproduct (%)** |
|---|---|---|---|---|---|---|---|
| **1** | HDAC6 SEC | 1 | 100 | 3 | 30 | 67 | |
| **2** | HDAC6 SEC | 1 | 300 | 47 | - | 53 | |
| **3** | HDAC6 SEC | 1 | 600 | 61 | - | 39 | |
| **4** | HDAC6 SEC | 0 | 100 | 100 | - | - | |
| **5** | HDAC6 SEC | 0 | 300 | 100 | - | - | |
| **6** | HDAC6 SEC | 0 | 600 | 100 | - | - | |
| **7** | HDAC10 SEC | 2 | 100 | - | 34 | 56 | 10 |
| **8** | HDAC10 SEC | 2 | 300 | - | 8 | 92 | |
| **9** | HDAC10 SEC | 2 | 600 | 31 | - | 69 | |
| **10** | HDAC10 SEC | 0 | 100 | 100 | - | - | |
| **11** | HDAC10 SEC | 0 | 300 | 100 | - | - | |
| **12** | HDAC10 SEC | 0 | 600 | 100 | - | - | |
| **13** | HDAC6 SEC | 10 | 100 | - | 96 | 4 | |
| **14** | HDAC6 SEC | 100 | 100 | - | 74 | 9 | 17 |
| **15** | HDAC10 SEC | 10 | 100 | - | 98 | 2 | |
| **16** | HDAC10 SEC | 100 | 100 | - | 70 | 7 | 23 |

**Single-Point HDAC6 Inhibitory Activity Assay in the Presence and Absence of TCEP**

Measurements were made using a custom fluorogenic peptide substrate Ac-RHKK(Ac)-AMC (Ac = acetyl, AMC = 7-amino-4-methylcoumarin). In a Corning 96-Well Solid Black plate, 25 µL of assay buffer [50 mM Tris-HCl (pH 8.0), 137 mM NaCl, 2.7 mM KCl, 1.0 mM $MgCl_2$] containing 1 µM HDAC6 were combined with 5 µL of SEC buffer containing 100 µM SelSA-2 or SAHA, with and without 1 mM TCEP. To initiate the reaction, 20 µL of assay buffer containing 375 µM substrate was added and the reaction was allowed to proceed for 30 min. To quench, 50 µL of developer solution (1 µM trypsin and 10 µM Tubastatin A in assay buffer) was added to the reaction mixture and allowed to equilibrate for 30 min. Fluorescence readings were measured on a Tecan Spark Multimode Microplate Reader (excitation = 360 nm, emission = 460 nm). All reactions were run with three technical replicates, including control experiments consisting of no enzyme, no substrate, and no inhibitor (Figure S2). Data were normalized from 100% to 0% using the average values of controls (100% = no inhibitor, 0% = no enzyme/no substrate).

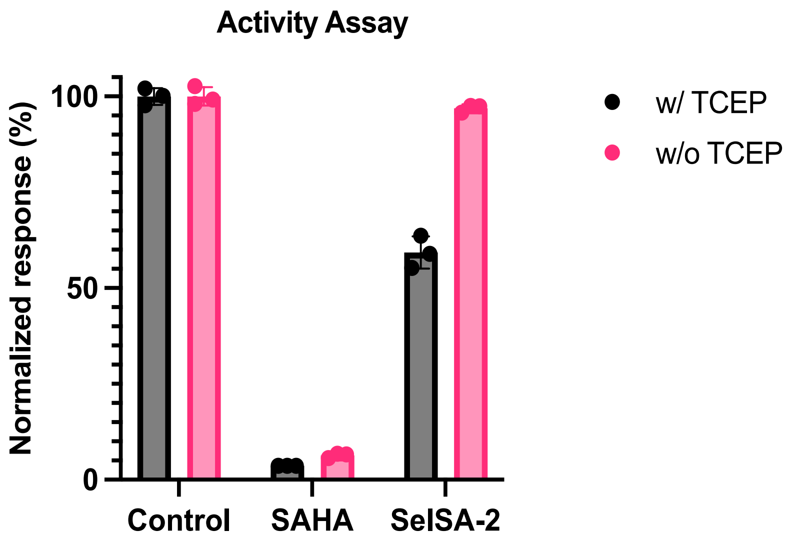


**Figure S2.** Fluorometric assays performed with HDAC6 in the presence and absence of reducing agent.
